# Amplicons and isolates: *Rhizobium* diversity in fields under conventional and organic management

**DOI:** 10.1101/2020.09.22.307934

**Authors:** Sara Moeskjær, Marni Tausen, Stig U. Andersen, J. Peter W. Young

## Abstract

**Background:** The influence of farming on plant, animal and microbial biodiversity has been carefully studied and much debated. Here, we compare an isolate-based study of 196 *Rhizobium* strains to amplicon-based MAUI-seq analysis of rhizobia from 17,000 white clover root nodules. We use these data to investigate the influence of soil properties, geographic distance, and field management on *Rhizobium* nodule populations.

**Results:** Overall, there was good agreement between the two approaches and the precise allele frequency estimates from the large-scale MAUI-seq amplicon data allowed detailed comparisons of rhizobium populations between individual plots and fields. A few specific chromosomal core-gene alleles were significantly correlated with soil clay content, and core-gene allele profiles became increasingly distinct with geographic distance. Field management was associated with striking differences in *Rhizobium* diversity, where organic fields showed significantly higher diversity levels than conventionally managed trials.

**Conclusions:** Our results indicate that MAUI-seq is suitable and robust for assessing nodule *Rhizobium* diversity. We further observe possible profound effects of field management on microbial diversity, which could impact plant health and productivity and warrant further investigation.

## Introduction

The interplay of plants and microorganisms in the soil has a multitude of beneficial functions in natural ecosystems, including protection against pathogens (Berendsen *et al*., 2012; Schlatter *et al*., 2017) and abiotic stress such as drought, uptake of nutrients like phosphate and nitrogen(Oldroyd *et al*., 2011; Gutjahr & Parniske, 2013), and growth promotion (Panke-Buisse *et al*., 2015).

Understanding the microbial variability within and between fields, and which factors influence the number or diversity of microbes, is necessary to understand how to best optimise or work with the microbiome. The increase in genetic diversity over distance for microorganisms has been shown at scales ranging from metres to 100 kilometres (Whitaker *et al*., 2003; Ramette & Tiedje, 2007). This effect can be explained by sampling a wide range of conditions or by isolation-by-distance, first described in aquatic *Sulfolobus*, and since then verified in multiple species (Whitaker *et al*., 2003; Rosselló-Mora *et al*., 2008; Vos & Velicer, 2008; Hahn *et al*., 2015).

Biological nitrogen fixation (BN)F occurs as a result of a mutualistic symbiosis between legumes and soil bacteria, commonly known as rhizobia. Rhizobia are harboured in specialised root structures, known as nodules. To confidently establish the level of diversity within nodule populations, the most common assessment method uses cultured bacteria isolated from nodules (Sbabou *et al*., 2016; Efrose *et al*., 2018; Stefan *et al*., 2018; Boivin *et al*., 2020; Cavassim *et al*., 2020). Isolate-based approaches rely on the culturability of the microbes and become very labour intensive if the desired number of isolates per site is high, though they have the advantage that isolates are available for evaluation as potential inoculants. For soil microbiome diversity studies, where many of the organisms cannot be cultured using traditional methods, high throughput amplicon sequencing (HTAS) is used to amplify sequences that distinguish microbial communities at different levels of resolution from environmental DNA samples in a cultivation-independent manner (Smalla *et al*., 2001; Costa *et al*., 2006). This method can be adapted for *Rhizobium* nodule or soil populations using multiplexed amplicons with unique molecular identifiers (MAUI-seq), as has been shown in recent publications (Fields *et al*., 2019; Boivin *et al*., 2020). How well diversity estimates from traditional isolate-based approaches compare to HTAS in evaluating the rhizobial diversity has not been explored in detail.

A number of studies have addressed the influence of land management on the soil microbiome by comparison with undisturbed soils such as native tropical forests and permanent grasslands (Palmer & Young, 2000; Mendes *et al*., 2015; Coller *et al*., 2019). Land management was found to have an impact on the alpha-diversity of fungal and bacterial microbial communities. Managed land shifted the balance towards more abundant fungal communities at the expense of bacterial diversity. Application of nitrogen was shown to reduce alpha-diversity of protist, bacterial, and fungal soil communities in maize fields, with possible detrimental effects on the soil microbiome (Zhao *et al*., 2019). Standard agricultural practices involve applying large amounts of N, both as synthetic and organic fertiliser, to ensure the yield and quality of the crop (Hansen *et al*., 2000), with possible detrimental effects on microbiome diversity.

Defining the way in which land management influences the soil microbiome is important when moving towards more sustainable agricultural practices. For organic farmers, where nitrogen input is limited, BNF is essential in fertilising the soil and providing high-yield crops with high protein content. The health and biodiversity of the soil microbiome can influence the yield of legumes through this symbiosis by affecting the available genotypes of rhizobia (Heath & Tiffin, 2009). The changes to the soil microbiome by management and fertiliser application could affect the nitrogen (N) availability in agricultural settings by limiting legume nodulation efficiency due to decreased abundance of rhizobia and other nodule-associated bacteria (Martínez-Hidalgo & Hirsch, 2017).

For some legumes grown in non-native soils, an effective symbiont is not naturally present. The solution to this issue has been to inoculate fields with the appropriate rhizobial symbiont for the legume crop. However, these inoculum rhizobia can be outcompeted by native rhizobia before the end of the growth season, even in soils with low levels of native rhizobia (Thies *et al*., 1991). Therefore, when growing legumes in native soils with a high concentration of rhizobia, where inoculation is an even less effective tool than in low-rhizobia soils, it is important to maintain a rich diversity of highly adapted microbes in the soil to ensure that an appropriate, effective symbiont partner is present for the legume crop of choice (Stajkovic-Srbinovic *et al*., 2012).

White clover (*Trifolium repens* L.) is an important agricultural crop in temperate climates used by both conventional and organic farmers primarily to improve forage quality by raising protein content in perennial grass pastures. Its symbiotic partner, *Rhizobium leguminosarum* (*Rl*), is a species complex comprising at least seven genetically distinct genospecies (gs) with limited gene flow between them (Kumar *et al*., 2015; Boivin *et al*., 2020; Cavassim *et al*., 2020). *Rl* can nodulate several legume species, and its specificity is determined by a group of symbiosis genes located on mobile plasmids. The population of *Rl* capable of establishing a symbiosis with clovers is the symbiovar *trifolii* (*Rlt*). Recently, methods investigating microbial intraspecies diversity in environmental samples have been developed. MAUI-seq, which is the method used in this study, relies on multiplexed amplicons tagged with unique molecular identifiers (UMIs) (Fields *et al*., 2019). UMIs allow filtering of erroneous reads (chimaeras and polymerase errors) using a ratio of how often a sequence is observed as a primary UMI sequence (the most abundant sequence tagged with a given UMI) or a secondary sequence (a less abundant sequence for a given UMI).

We have previously characterised a set of genomes from 196 *Rlt* isolates from pink white-clover nodules from three clover field trial sites in northern Europe, and 50 organic fields from Jutland, Denmark (Cavassim *et al*., 2020). These 196 genomes were distributed throughout five of the seven known genospecies in *Rl*, with the majority belonging to gsC. Here, the *Rlt* nodule populations in the same fields were characterised using the MAUI-seq method (Fields *et al*., 2019) by amplifying two core genes (*rpoB* and *recA*) and two accessory genes (*nodA* and *nodD*) important in establishing symbiosis. We compare HTAS with the traditional isolate-based approach in evaluating the intraspecies *Rlt* diversity in white-clover nodule populations in field trials and organic fields, to investigate the allelic diversity at each site in greater depth.

While isolates potentially provide the full genome information and allow assessment of whole genome differences between strains, sample sizes are necessarily limited. In recent studies, the numbers of isolates ranged from 73 to 212 ((Efrose *et al*., 2018) n=73; (Stefan *et al*., 2018) n=86; (Cavassim *et al*., 2020) n=196; (Boivin *et al*., 2020) n=210; (Sbabou *et al*., 2016) n=212). Our previous study characterised isolates from 196 nodules in detail and facilitated in-depth population genomics analysis and the discovery of movement of symbiosis genes between genospecies on a promiscuous plasmid (Cavassim *et al*., 2020). Using MAUI-seq, we were able to process and study the nodule population on a much larger scale, obtaining sequencing data of amplicons from 17,000 nodules.

We found that isolate-based and MAUI-seq diversity assessment were similar in terms of genospecies abundance and that all highly abundant sequences overlapped. MAUI-seq identified more rare alleles for all amplicons except *recA*. We concluded that the diversity observed was robustly determined by both methods, and a small set of chromosomal core genes and plasmid-borne accessory nod genes were significantly correlated with differences in soil clay and silt content. Core genes were affected by isolation by distance to a greater extent than plasmid-borne symbiosis genes in a set of samples from organic fields in Jutland. When comparing genetic diversity in nodule populations from fields under different management, samples from organic fields had significantly higher genetic diversity than fields used for conventional clover breeding trials, indicating that biodiversity of clover symbionts is affected by field management.

## Results

We previously characterised 196 *Rlt* genomes isolated from pink nodules collected from 40 plots in three clover breeding trial sites at Rennes in France (F), Didbrook in England (UK), and Store Heddinge in Denmark (DK), as well as from 50 fields on Danish organic farms (DKO) (**Figure 1**). The strains were distributed throughout five of the seven known genospecies for *Rl*, with some genospecies being highly site-specific. gsC was the dominant genospecies at all sites except the UK trial site, where only gsB was present. Conversely, gsB was only found at the UK site.

**Figure 1.**
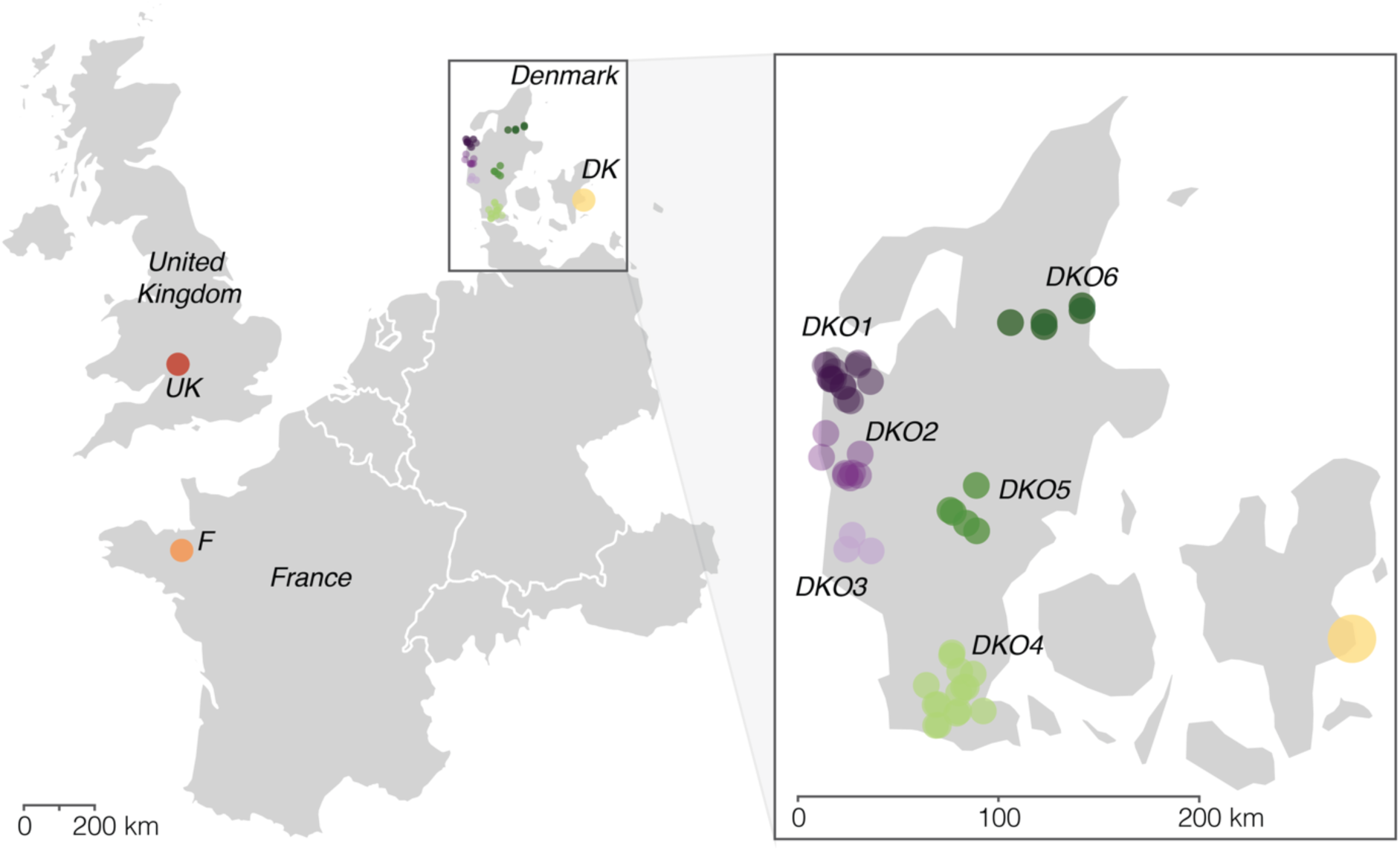
Sampling sites. Clover breeding trial sites in Rennes (F), Didbrook (UK), Store Heddinge (DK). Each dot represents 40 samples. Organic fields sampled in Jutland (DKO1-6). Each dot represents one sample. The total number of sample sites is 170 (UK=40, F=40, DK=40, DKO1=14, DKO2=8, DKO3=3, DKO4=15, DKO5=5, DKO6=5).

To investigate whether this set of 196 genomes is representative of the *Rlt* diversity at the sampled sites we collected nodules from the plots within the trials sites and the organic fields, leaving us with a total of 170 samples of nitrogen-fixing white clover root nodules from European field sites (**Figure 1, Figure S1**). Using the MAUI-seq method (Fields *et al*., 2019), 100 pooled nodules per sample yielded genotype frequencies of two core (*rpoB* and *recA*) and two accessory (*nodA* and *nodD*) genes in 170 samples. After filtering for samples with missing data or low UMI count, 105 *rpoB* samples, 153 *recA* samples, 129 *nodA* samples, and 130 *nodD* samples were used for downstream analysis.

### Geographically distinct sites display a site-specific set of nodule *Rhizobium* alleles

*Rhizobium leguminosarum* is a species complex consisting of multiple genospecies that have been shown to co-exist in a field setting (Kumar *et al*., 2015; Boivin *et al*., 2020; Cavassim *et al*., 2020). *Rlt* core genes show little sign of introgression between genospecies, and phylogenies of individual core genes therefore most often follow the overall genospecies phylogenetic tree (Stefan *et al*., 2018; Cavassim *et al*., 2020). A phylogenetic analysis of amplicons from the chromosomal core genes *rpoB* and *recA* showed that the sampled bacteria from nodules are distributed throughout the five main genospecies clades previously identified from isolates originating from these exact fields (Cavassim *et al*., 2020) (**Figure 2A**-**D**). For the core genes *rpoB* and *recA*, the majority of the alleles identified by MAUI-seq were also recovered in the isolates, while some additional alleles were found only in a small number of isolates, particularly for *recA* (**Table 1** and **Figure 2A**-**D**). Of these sequences, most were actually present in the MAUI-seq dataset, but were under the cumulative abundance threshold that we used. For the other three genes, MAUI-seq recovered more alleles than the isolates.

**Figure 2.**
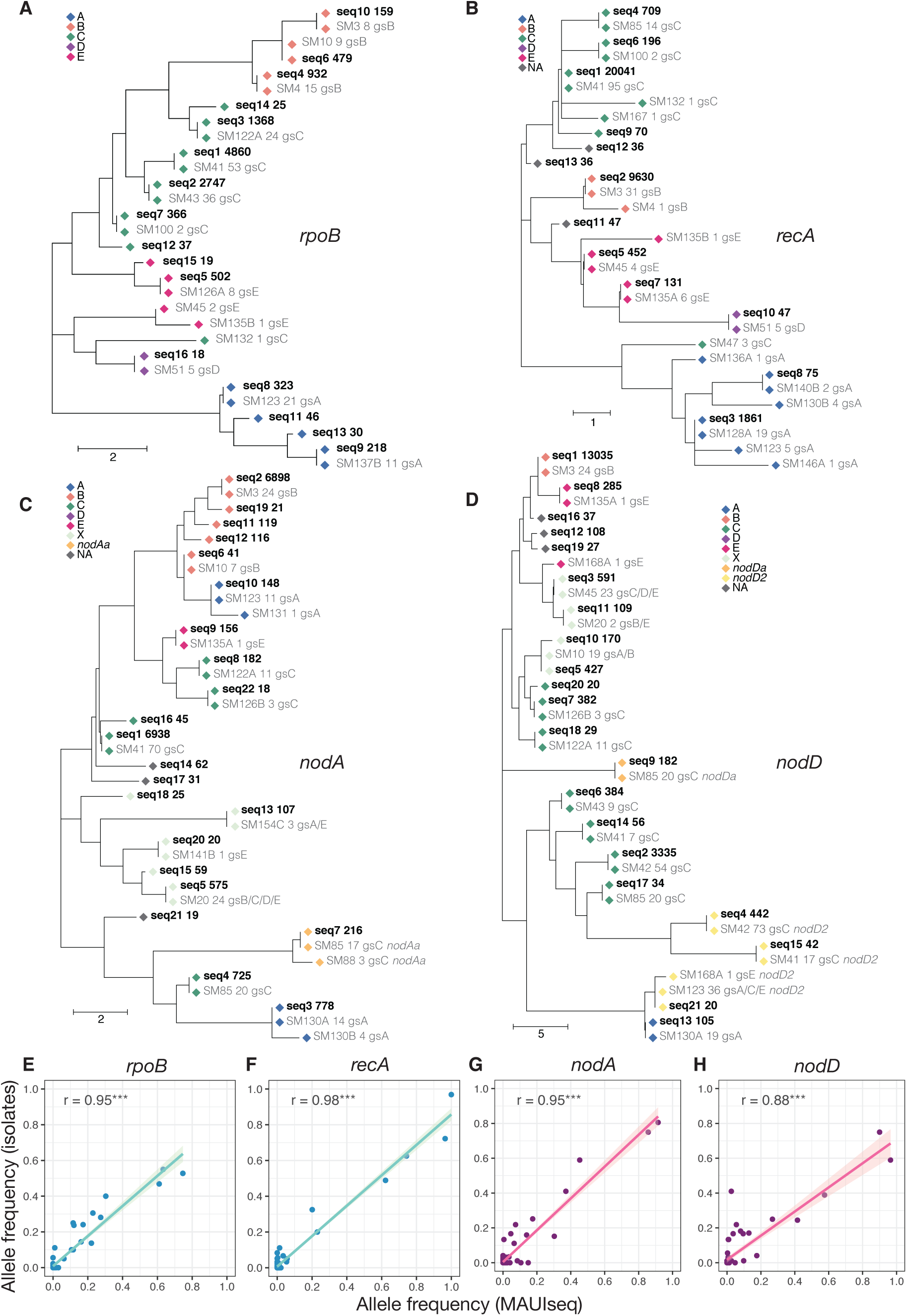
**A**-**D**: Phylogenetic trees of all alleles found in isolates or amplified from nodules.. MAUI-seq amplicons (black, bold) for each gene (nomenclature: seq - abundance rank - primary_UMI_count) and sequences (grey, light) from representative isolates (nomenclature: strainID - number of alleles in 196 genomes – genospecies) (Cavassim *et al*., 2020) have been included. The scale is in the number of nucleotide differences. Core genes (*rpoB* and *recA*) are assigned to the genospecies A-E (Kumar *et al*., 2015). Nod genes (*nodA* and *nodD*) are assigned to a genospecies if possible, or to a clade of introgressing genes labelled X (Cavassim *et al*., 2020). If an amplicon could not clearly be assigned to a clade it is marked as NA. **A:** *rpoB*. **B:** *recA*. **C:** *nodA*. **D:** *nodD*. **E**-**H**: Relative allele abundance for individual genes within sites (DK, F, UK, DKO) for the two different methods. Each point represents an allele that was found in the isolates and/or the MAUI-seq data. For each location (DK, F, UK, DKO), the frequency among isolates is plotted against the average frequency of the same allele in the MAUI-seq samples. In the case of DKO, where the number of isolates per field varied from 1 to 3, each field was weighted equally.

**Table 1.** Number of alleles identified both by MAUI-seq and in the 196 *Rlt* isolates (Cavassim *et al*., 2020), and those occurring exclusively in one of the datasets. Numbers in parentheses denote how many of the **196 *Rlt* solates only** sequences were recovered by MAUI-seq at a lower cumulative abundance threshold.

|  | *rpoB* | *recA* | *nodA* | *nodD* |
| --- | --- | --- | --- | --- |
| **Sequences in both datasets** | 11 | 9 | 12 | 12 |
| **MAUI-seq only** | 5 | 4 | 9 | 5 |
| **196 *Rlt* isolates only** | 3 (1) | 9 (6) | 2 (1) | 0 |

The accessory genes *nodA* and *nodD* belong to a group of co-located genes, known as the sym gene cluster, that are essential for initiating and maintaining an effective symbiotic relationship with legumes. The phylogeny of the accessory gene pool has previously been shown to often be incongruent with the core genes (Tian *et al*., 2010; van Cauwenberghe *et al*., 2014; Andrews *et al*., 2018; Efrose *et al*., 2018; Cavassim *et al*., 2020). This cluster is usually located on a conjugative plasmid in the *Rl* species complex (Kumar *et al*., 2015; Boivin *et al*., 2020; Cavassim *et al*., 2020). Occasionally, regions of the cluster are duplicated in the rhizobial genome and, due to the promiscuous nature of conjugative plasmids, they can cross genospecies boundaries (Cavassim *et al*., 2020). Using the set of 196 characterised *Rlt* isolates from the same sampling sites, we evaluated the level of duplication of *nod* genes to remove potential paralogs. In addition to the full nod gene region (*nodXNMLEFDABCIJ*), a partial set of nod genes (*nodDABCIJT*) is present in some of the *Rlt* isolates. nodAseq7 and nodDseq9 occurred only as secondary sequences in this partial nod region and were designated as *nodAa* and *nodDa*, respectively (**Figure 2C-D**). A third type of *nodD* (*nodD2*) was observed in some genomes flanked by transposases and no other nod genes (Kelly *et al*., 2018; Ferguson *et al*., 2020). Three *nodD* amplicons belong to this group. These five paralogous sequences were removed from all downstream analysis to avoid inflating the estimates of overall diversity. All 12 *nodD* alleles seen in the genomes were recovered by MAUI-seq, plus an additional 5 alleles. MAUI-seq detected 12 of the 14 *nodA* alleles seen in genomes, but found an additional 9 alleles (**Table 1** and **Figure 1**). All of the abundant sequences with frequency > 0.15 have an exact match in the 196 *Rlt* genomes, and the allele frequencies are highly correlated between the two datasets (**Figure 2E**-**H**). The sequences identified only by MAUI-seq are of low abundance, but appear to be genuine sequences (**Figure 2A**-**D** and **Figure 3**). Likewise, the sequences in the 196 genomes not found by MAUI-seq are only present in a small number of isolates and at low frequencies; 8 out of the 13 sequences are only found in a single isolate (**Figure 2**).

**Figure 3.**
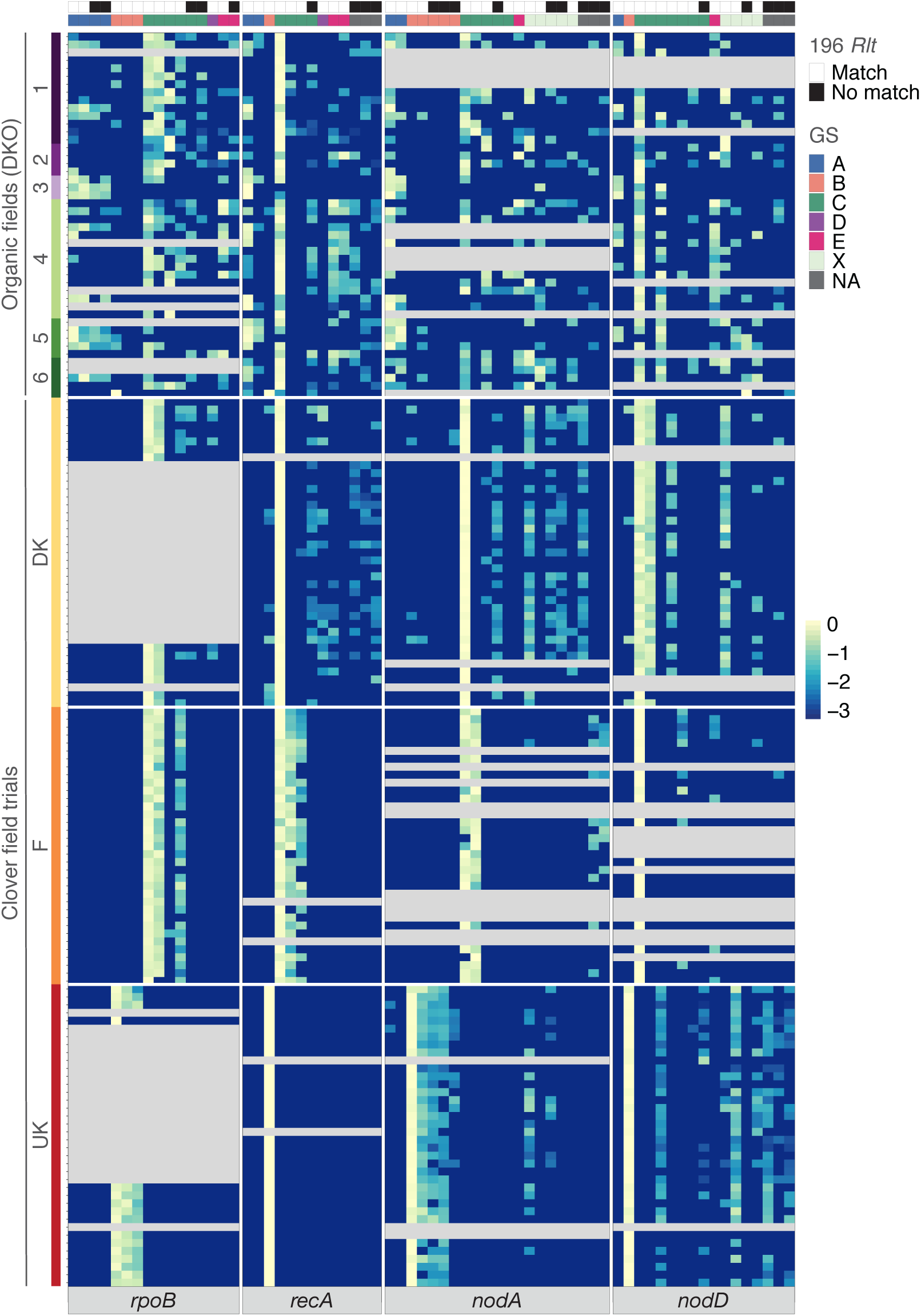
Heat map of relative amplicon frequency for individual genes. Samples with UMI count under 10 for an individual gene are in grey and are excluded from all analyses. The normalisation is done for each gene individually. Management regime and grouping of the samples are indicated to the left of the heat map. Sequences that have an exact match in the 196 *Rlt* genomes (196 *Rlt*), and the genospecies clade (GS) of each allele, are indicated by coloured bars above the heat map. (nSamples_rpoB: 105, nSamples_recA: 153, nSamples_nodA: 129, nSamples_nodD: 130). Abundance scale is log_10_(frequency).

Principal component analysis of the amplicons from individual genes (**Figure S2**), revealed that different loci have different levels of resolution. *recA* separated the French samples well from all other locations, whereas the UK samples were clearly separated from the other two field trial sites for all four loci. The high level of diversity among and within DKO samples made it difficult to distinguish them from the F and DK samples for most amplicons.

Each breeding trial site (DK, F, and UK) showed a distinct set of amplicons, despite the nodules from each site being sampled from the same F2 clover families from the same seedstock and being under identical management (**Table S1, Figure 3**). The samples from the trial sites were relatively uniform within each site, and each sample had a low number of total observed amplicons, whereas the DKO samples appeared less homogeneous within each sample.

### Genetic differentiation is correlated with spatial separation

To assess the biogeographic patterns of nodule populations, we calculated the hierarchical *F*_ST_ of samples at different population levels. The top level was management, where we compared organic fields (DKO) versus field trial sites, whereas plot and field were the lowest levels for the field trials and DKO samples, respectively (**Figure 1** and **Figure S1**). *S*ignificance was tested by permutation. For example, when comparing organic (DKO) versus conventional (field trial) management, samples were moved from one management to the other to check if this generally resulted in a lower *F*_ST_. For each management type, we then tested for the effect of country for the field trials or groupings for the DKO samples. We observed no differentiation between the conventionally managed sites (DK+F+UK) and the organic DKO population. The reason is that there is no overlap in *Rhizobium* populations between the UK and DK+F trial sites. Therefore the difference between the three trial sites is as high as the difference between trial sites and organic fields. To test the effect of groupings and field/plots on the differentiation, we then analysed field trial sites (DK, F, and UK) and the organic fields (DKO) separately. For the field trial subset, country (DK, F, and UK), had a significant effect (**Table 2**). The block design and clover genotype (**Figure S1**) did not have any effect on *F*_ST_ with the exception of *nodA*, where there was a small but highly significant effect. Permuting the plots within the individual field trial sites showed that the plots are also significantly associated with *Rhizobium* population differentiation. Furthermore, we added the block design and clover genotype level to the test (**Figure S1**) and found that both block and clover genotype had a small but significant effect on *nodA* differentiation when samples were permuted within the same country (**Table 2**).

**Table 2.** Hierarchical *F*_ST_ estimates for levels within the field trial subset and the DKO grouping subset. Levels of sampling are: **country level** (**Figure 1**, DK, F, and UK), **grouping level** (**Figure 1**, DKO1, DKO2, DKO3, DKO4, DKO5, DKO6), and **field/plot level** (individual samples from fields within groupings and plots within field trials, **Figure 1** and **Figure S1**). **Block level** and **clover genotype level** (**Figure S1**) are also included for the field trial subset. Numbers show the *F*_ST_ estimates of each level out of the total variance (e.g. *F*country/total) and within outer levels (e.g. effect of block level within each country: *F*block/country). Statistically significant values compared to 1000 random permutations are indicated with asterisks: *p<0.05, **p<0.01, and ***p<0.001.

| Field trial sites |  |  |  |  |  |
| --- | --- | --- | --- | --- | --- |
|  |  | *rpoB* | *recA* | *nodA* | *nodD* |
| **Country** | $F_{country/total}$ | 0.847*** | 0.892*** | 0.736*** | 0.772*** |
| | $F_{block/country}$ | 0 | 0 | 0.023*** | 0 |
| | $F_{clover/country}$ | 0 | 0 | 0.018*** | 0 |
| | $F_{plot/country}$ | 0.121*** | 0.139*** | 0.237*** | 0.128*** |
| DKO |  |  |  |  |  |
|  |  | *rpoB* | *recA* | *nodA* | *nodD* |
| **Grouping** | $F_{grouping/total}$ | 0.198** | 0.202** | 0.025 | 0.001 |
| | $F_{field/grouping}$ | 0.569*** | 0.504*** | 0.465*** | 0.459*** |
| **Field** | $F_{field/total}$ | 0.654*** | 0.604** | 0.478* | 0.462*** |

In the DKO subset, the grouping (DKO1, DKO2, DKO3, DKO4, DKO5, DKO6), based on geographic proximity (**Figure 1**), had a significant effect on differentiation for the core genes, *recA* and *rpoB*, but not for the accessory genes, *nodA* and *nodD* (**Table 2**). Field had a significant effect for all four genes within the DKO subset. We concluded that the grouping level was a valid way to cluster the samples for the cores genes, and used the DKO subset to calculate pairwise *F*_ST_ between the six DKO clusters (**Figure 1** and **Figure 4**). The fields differ in geographic placement, while the clover genotypes and the field management were similar between sites, allowing us to explore the geographic differences in nodule population within a homogeneous set of managements (**Table S1**).

**Figure 4.**
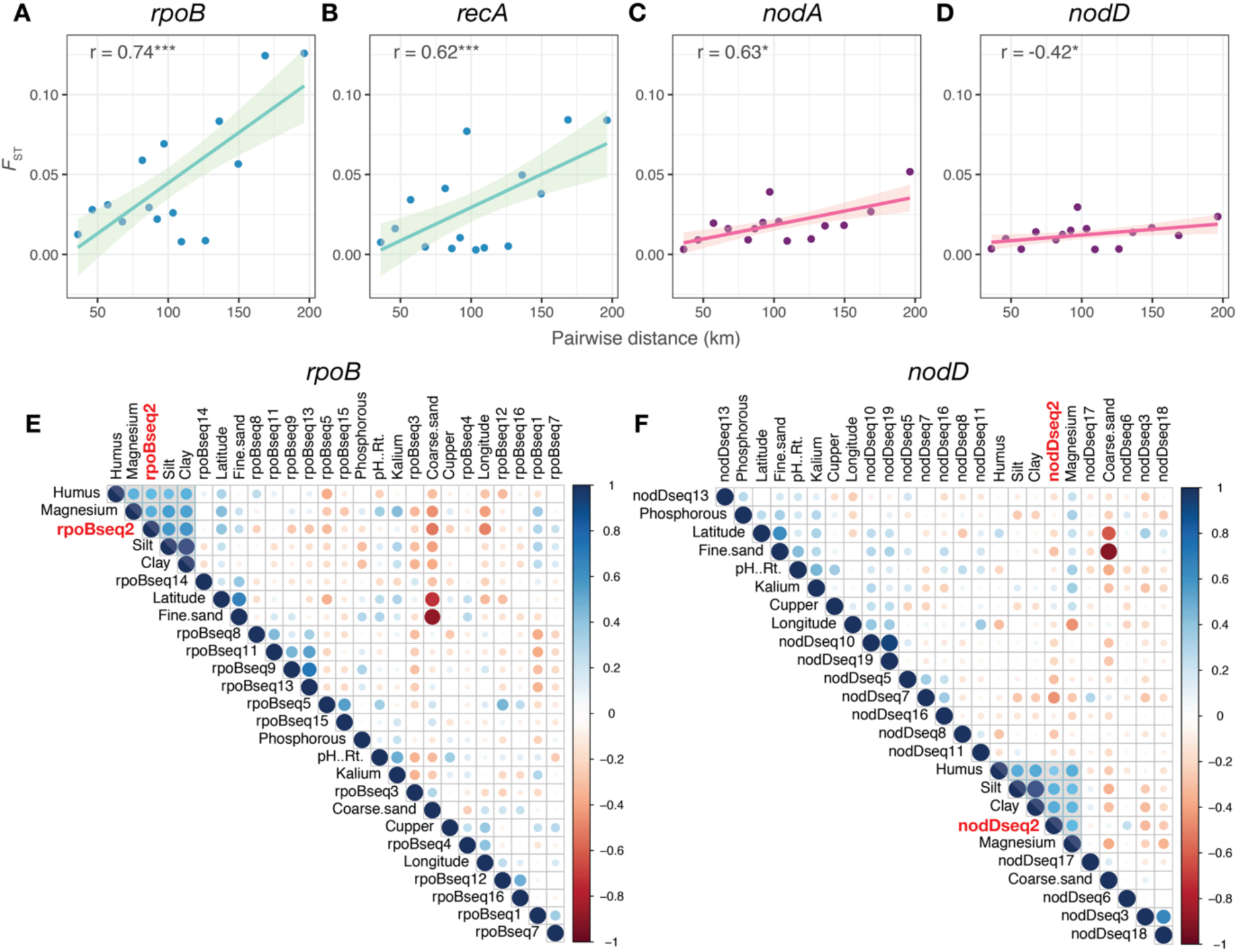
Correlation between increase in genetic diversity (*F*_ST_) and geographic distances for pairwise comparisons between DKO samples. p-values are indicated by asterisks; *p<0.05, **p<0.01, and ***p< 0.001. **A-D:** Pairwise *F*_ST_ between DKO clusters. **E-F:** Correlations between normalised allele frequencies and soil chemical properties per gene for the DKO subset for a core gene (*rpoB*, **E**) and an accessory gene (*nodD*, **F**). The cluster of high correlations including clay and silt are highlighted in grey. Alleles within the cluster are highlighted in red.

Pairwise *F*_ST_ between the DKO clusters displays a significant correlation between *F*_ST_ and geographic distance. The overall *F*_ST_ is highest for the two core genes, and the correlation is highest and most significant for *rpoB* (**Figure 4**). The level of genetic differentiation between populations increased with distance, with the effect being most pronounced for chromosomal core genes, indicating that the core gene population composition is related to geographic origin. Since the samples are not independent, we performed a Mantel test with 5000 replicates to test if the correlation between *F*_ST_ and geographical distance is significantly different from randomised datasets. Three genes, *rpoB, recA*, and *nodA*, have a significant correlation between genetic differentiation and geographical distance (*rpoB*: R^2^=0.737, p-value=0.002, slope=6.310e-07; *recA*: R^2^=0.617, p-value=0.019, slope=4.156e-07; *nodA*: R^2^=0.632, p-value=0.047, slope=1.748e-07), but *F*_ST_ and geographical distance are not significantly correlated for *nodD* (*nodD*: R^2^=0.420, p-value=0.115, slope=7.058e-08).

### Limited allele correlation with soil chemical properties

Adaptations to ecological niches require different sets of genes. Genetic differentiation in soil microbial communities is therefore often linked with the chemical and physical composition and pH of the soil. To test whether the correlation between *F*_ST_ and geographical distance was due to geographically linked differences in soil chemical composition, we tested the correlations between allele frequency and soil traits from the fields where the samples were collected.

Several clusters of strong correlations between allele frequency and soil chemical and physical properties were observed for the full dataset (**Figure S4A-D**). The high clay content in the UK field trial site and the unique set of gsB alleles observed in these samples drove this clustering (**Table S1, Figure 2**). Since none of the UK gsB alleles were observed in any other samples, no conclusions can be drawn as to whether the gsB dominance is due to an increased fitness in clay, or to geographical influence.

To test a more homogeneous set of samples, we focused on the DKO subset, which had a broad range of values for all soil chemical properties and no extreme values that could be confounded with rare alleles. Two core gene alleles, rpoBseq2 (correlation with clay = 0.6141, p-value = 5.03e-05; silt: correlation = 0.5877, p-value = 0.0001) and recAseq4, and one common *nod* allele (nodDseq2) were highly correlated with silt and clay content, (**Figure 4E** and **4F, Figure S4**). The *recA* allele was very rare, and only occurred in four samples, whereof two had a high silt and clay content, driving the correlation signal. rpoBseq2 and nodDseq2 were both correlated with silt and clay. Both alleles were assigned to genospecies C (**Figure S2**), and had a correlation of 0.525 (p-value = 0.0029). To investigate whether these two alleles co-occur within the *Rlt* isolates, we BLASTed them against the 196 whole genome sequences. The *rpoB* allele was present in 36 of the genomes, whereas the *nodD* allele was present in 55 genomes. The alleles co-occurred in 30 strains, most of which were isolated in fields or field trials sites with a high clay/silt content, suggesting the genomic architecture of these strains might confer some increased fitness in clay/silt rich soils. The majority of strong correlations observed were between alleles, meaning some strains tend to co-occur, or between soil chemical properties that are correlated (such as silt and clay) or mutually exclusive (such as coarse sand and fine sand). Since no alleles or soil chemical properties are highly correlated with latitude or longitude, the *F*_ST_ correlation with geographical distance (**Figure 4**) does not seem to be driven by differences in soil chemistry or composition.

### Bacterial richness within samples is higher for fields under organic management

To assess the effect of field management on *Rhizobium* diversity, we analysed the genospecies composition of each individual sample. The amplicons from all four genes were assigned to genospecies A-E (**Figure S2**) (Kumar *et al*., 2015). *nodA* and *nodD* amplicons were assigned as X if they were within an introgressing clade of plasmid-borne sym genes (**Figure S2C-D**) (Cavassim *et al*., 2020). The genospecies composition was plotted for each individual sample of 100 nodules (**Figure 5**). At two of the trial sites (F and UK), only core genes from a single genospecies were detected, gsC and gsB, respectively (**Figure 5A** and **B**), though several alleles of the same genospecies were present (**Figures 2 and 5**). The DK trial site had low levels of gsE, gsD, and gsB, while the dominant clade was gsC. For the DKO samples, core genes belonging to all genospecies clades were found, and most samples contained several genospecies at intermediate frequencies. The raw UMI count was not higher for the DKO samples, indicating that the increase in observed genospecies is not due to sequencing bias (**Figure S3**). The diversity of core genes, *rpoB* and *recA*, for the F and UK trial sites matched the genospecies distribution of isolates from these fields. For the DK site, low levels of core and accessory gsB amplicons were found, although no gsB strains were isolated from here. The local gs distribution matches the observations from our previous study of isolates from these sites (Cavassim *et al*., 2020), indicating that the sampling of nodule isolates from fields can yield a good approximation of the actual rhizobium population present in the nodules (**Figure 5**).

**Figure 5.**
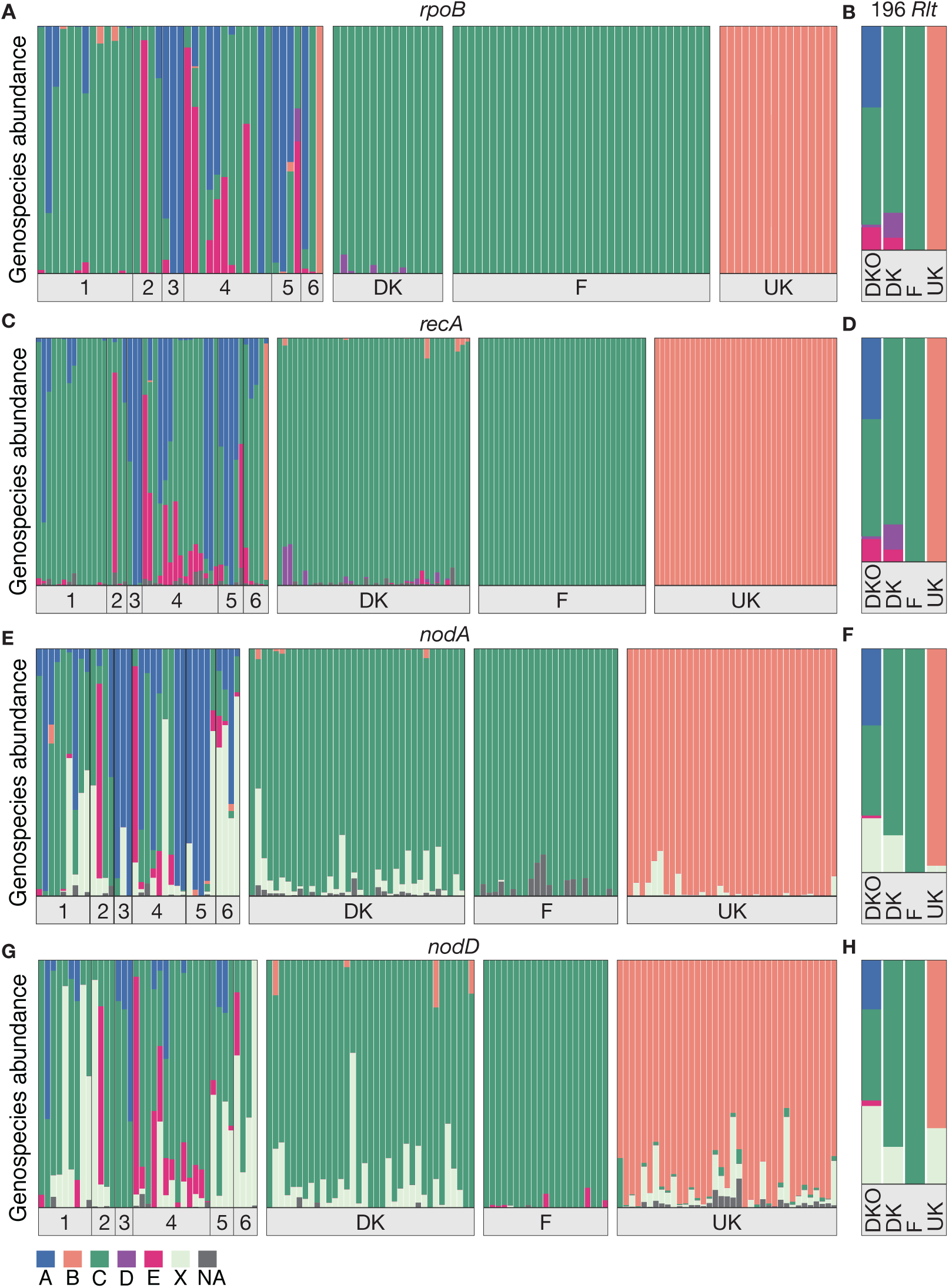
Genospecies composition of *Rlt* from nodules. **A, C, E**, and **G**: Genospecies composition of each individual sample for each gene (**A**: *rpoB*, **C**: *recA*, **E**: *nodA*, and **G**: *nodD*). The DKO groupings are labelled by their respective number (DKO1=1). Core genes (*rpoB* and *recA*) are assigned to the genospecies A-E (Kumar *et al*., 2015). Nod genes (*nodA* and *nodD*) are assigned to a genospecies if possible, or to a clade of introgressing genes labelled X (Cavassim *et al*., 2020). If an amplicon could not clearly be assigned to a clade it is marked as NA. **B, D, F**, and **H**: Genospecies composition based on individual genes of isolates from DKO fields (n=88), DK (n=36), F (n=40), UK (n=32) for *rpoB, recA, nodA*, and *nodD*, respectively (Cavassim *et al*., 2020).

Overall, more genetic diversity was observed within the samples for the accessory genes than for core genes for most sampling sites, predominantly due to the introgressing clade, X. Furthermore, most samples from the three trial sites contained low levels of *nod* gene amplicons that were not assigned to the core gene genospecies observed in the samples (**Figure 5E** and **G**). Most of these belong to the introgressing clade, X, which is consistent with previous observations on large *Rlt* populations (Cavassim *et al*., 2020).

Some nod gene amplicons do not match the core gene composition. The French samples are exclusively gsC for the core genes, *rpoB* and *recA*, whereas the UK samples are exclusively gsB (**Figure 5A** and **5C**). The accessory genes, *nodA* and *nodD*, include sequences designated as gsE and gsC for the French and UK samples, respectively, (**Figure 5E** and **5G**), based on the genospecies designation from the 196 *Rlt* isolates. This indicates that these sym gene alleles introgress between genospecies, but this introgression event was not captured by the isolates (Cavassim *et al*., 2020).

To investigate whether the microbial diversity was higher for DKO than for the field trial samples, we calculated the nucleotide diversities (π). Comparing the nucleotide diversity of the individual DKO groupings to the three separate field trial sites, most DKO groupings had a significantly higher π than the DK, F, and UK trial sites (**Figure 6**). Grouping the samples by field management revealed that overall the organic DKO samples have significantly more diversity than the field trial samples for all four amplicons (*rpoB*: p-value=3.007e-06, *recA*: p-value=8.601e-07, *nodA*: p-value=1.916e-09, *nodD*: p-value=2.875e-08) (**Figure S5**). For our samples, this increase in nucleotide diversity within nodule populations indicates that the microbial diversity of the *Rhizobium* population is higher in fields under organic management than in field trial sites under conventional management.

**Figure 6.**
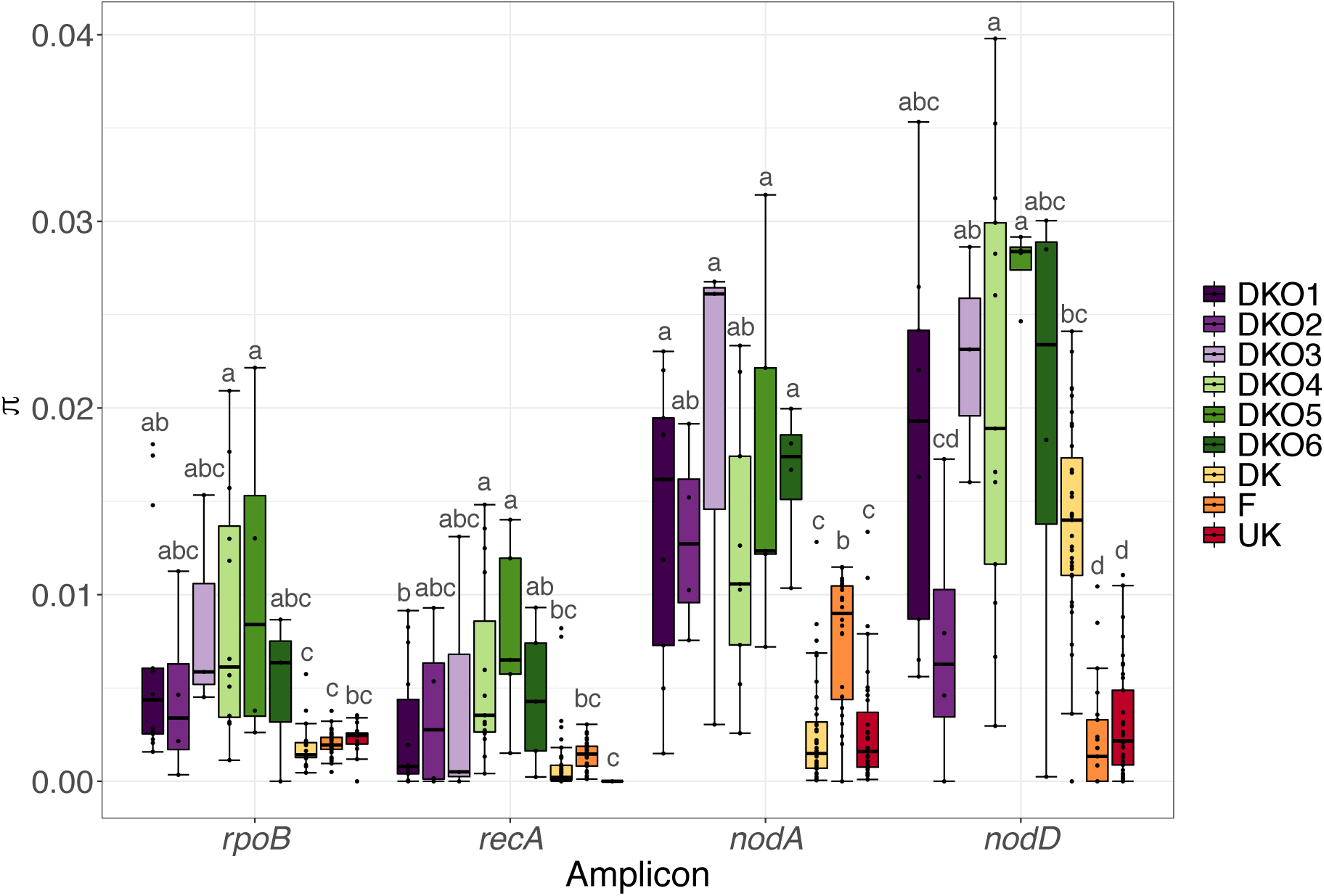
Nucleotide diversity within populations for each gene. π for individual samples within the DKO groupings and DK, F, and UK field trials. Dots illustrate the π value for each individual sample. Bars represent the first and third quartiles, with the solid line denoting the median. Whiskers correspond to the 1.5 * interquartile range. p-values were calculated for each individual gene using ANOVA followed by Tukey’s *post hoc* testing. Groupings indicated by the same letter were not significantly different at p<0.05.

## Discussion

The increased focus on sustainable agricultural practices has increased the need for knowledge on the impact of land management on biodiversity of all living things from mammals to fungi and bacteria. For legume crops, a healthy soil microbiome, and especially the availability of nitrogen-fixing rhizobial symbionts capable of establishing an effective symbiosis, plays an important role in establishing a high-yielding agricultural practice.

Most studies to assess field bacterial diversity have focused on *Rhizobium* isolates from *Trifolium* and *Vicia* nodules (Sbabou *et al*., 2016; Efrose *et al*., 2018; Stefan *et al*., 2018; Cavassim *et al*., 2020). Here, we present an in-depth study of clover field trials and organic fields where strains have previously been isolated (Cavassim *et al*., 2020), using the MAUI-seq method (Fields *et al*., 2019) on nodule populations to investigate the reported level of diversity within the *R. leguminosarum* community found in white clover nodules when using a HTAS approach compared to a more traditional isolate-based approach. It has been estimated that 99% or more of environmental prokaryotes cannot be cultured using classical methods in the laboratory. These obvious limitations are a challenge for many bacterial species, and HTAS is commonly used to investigate communities without the need for cultivation (Costa *et al*., 2006). *R. leguminosarum* is easily cultured from nodules in the laboratory, but it is slow growing and requires several rounds of single colony isolation before it can be separated from faster growing bacteria present in and around the nodule. Studying large numbers of nodules therefore requires both time and resources. HTAS offers the advantage that it allows for screening a large number of samples with many organisms per sample in a fast and efficient manner. We have adapted this method using MAUI-seq to study the intraspecies diversity of *R. leguminosarum* in nodules, to allow screening of large nodule populations from many sampling sites. While amplicon studies are ideal for sampling large numbers of samples (here, nodules), good quality sequences of the organism of choice are necessary to design specific, yet effective primers for any non-standard amplicon. The 196 sequenced *Rlt* isolates have formed the basis of the phylogenetic analysis and thereby the introgression analysis of this study. Here, we use amplicon sequencing to check that the isolates are representative of the nodule population as a whole.

There was good agreement between the two methods in estimating the allele frequency and genospecies composition of the nodule populations, though more rare alleles were observed in the MAUI-seq data probably due to the deeper sampling; 85 times more nodules were sampled using MAUI-seq compared to the isolate-based study. Many amplicon-based studies have assessed enrichment of many bacterial species in the rhizosphere or other soil compartments compared to bulk soil or between managements (Baudoin *et al*., 2003; Costa *et al*., 2006; Schreiter *et al*., 2014; Coller *et al*., 2019). Similarly, a modified version of MAUI-seq has been used to investigate the *Rhizobium* population in soil and how it differs from nodule populations from *Vicia* and *Trifolium* host plants of *Rhizobium* (Boivin *et al*., 2020). Cultivating slow-growing bacteria from soil can be very difficult, and amplicon sequencing enables studies of populations in bulk soil that would otherwise have been unfeasible.

Using the broadly sampled set of nodule amplicons allowed us to make comparisons between nodule populations in soils under different managements and with different soil chemical and physical properties. Genetic differentiation for accessory genes is less correlated with distance than for core genes, perhaps reflecting differences in local adaptation. It seems likely that *nodD* and *nodA* are not adapted to local differences, but are adapted to the available symbiotic partner (here, white clover). The movement of symbiosis genes may be less restricted since they are located on conjugative plasmids (Cavassim *et al*., 2020). Mobility of accessory genes located on integrative conjugative elements (ICEs) has previously been hypothesised to lead to lower genetic differentiation over distance than core genes located on less mobile parts of the chromosome (Hoetzinger *et al*., 2017). In our case, the accessory symbiosis genes, *nodA* and *nodD*, are located on conjugative plasmids shown to cross genospecies boundaries, whereas there is little to no recombination between core genes (Cavassim *et al*., 2020). Isolation-by-distance has been shown to be more pronounced for the core genome for aquatic microbes, whereas the location of accessory genes on plasmids or on genomic islands renders them more mobile and more dispersed in the environment (Hoetzinger *et al*., 2017), which might also be true for bacterial populations in soil. In-depth analysis of allele frequencies in each sample is possible, due to the number of nodules sampled when using amplicon sequencing. Using only the isolate dataset, we would be limited to a few isolates per field/plot, and hence imprecise estimates of diversity.

Correlations between genetic differentiation and distance between populations are often confounded by differences in soil physical and chemical properties. We tested the correlations between soil chemical and physical properties, geospatial placement, and amplicon abundance. No significant correlations were observed between latitude/longitude and soil properties. The observed positive correlation between genetic differentiation and spatial separation is therefore unlikely to be due to soil chemical or physical properties. The UK site, which had a uniform and unique composition of only gsB, has the highest phosphorus (P) content. P content has been shown to be correlated with rhizobium population size in a previous study, but no genotyping was done (Wakelin *et al*., 2018). The high nutrient content might enhance the fitness difference between fast- and slow-growing *Rlt* strains (Leff *et al*., 2015), thereby driving the population differences between high and low P sites.

The level of nucleotide diversity (**π**) observed within each nodule population sample was significantly higher for samples from fields under organic management than from fields used for clover breeding trials. This might reflect a lower diversity of *Rlt* in the soil at the clover breeding trial sites, possibly due to an increase in nitrogen application (Zhao *et al*., 2019). The DK, F, and UK sites are very different in soil composition, making it less likely that the lower *Rlt* genetic diversity is due to a soil-management interaction than to a general effect of field management. The *Rhizobium* populations at the three sites have distinct sets of alleles, suggesting that the management does not select for a specific set of *Rlt*, but rather enriches already dominant or highly adapted strains, specific for each site. A more diverse collection of clover genotypes was grown at the DK, F, and UK sites than for DKO fields, so the reduction in diversity is unlikely to be an effect of clover genotype selection. The varied genospecies distribution of isolates from the DKO fields hinted at a higher diversity in these fields compared to the field trial sites, but the differences in sampling distance between DKO isolates (up to 200km) and isolates from individual field trials (<200m) impeded a detailed investigation. For the amplicon-based study, we collected 100 nodules from each field, which allowed us to treat each field as an individual data point in the diversity analysis. This revealed the striking differences in nucleotide diversity between fields under different management regimes.

A study of spatial variation of *Rhizobium* symbiotic performance used a field sampling layout to test the effect at different spatial scales (Wakelin *et al*., 2018). Similar study designs, with neighbouring fields managed in different ways, would be appropriate for a more in-depth assessment of the effect of management on *Rhizobium* populations while disentangling it from geographical and soil chemical variation. Our results, in combination with previous studies, provide an indication that there could be substantial effects of field management on *Rhizobium* diversity and should motivate further studies on the effect of field management on soil microbial diversity at the level of individual species. We show that HTAS in the form of MAUI-seq on pooled nodules is an efficient method for estimating *Rhizobium* diversity in nodules, and a previous study has shown that the method can also be applied to soil samples (Boivin *et al*., 2020). There was good agreement between the alleles detected by amplicon sequencing and those found in isolates cultured from nodules at the same sites, and MAUI-seq can provide more detailed estimates of allele frequencies without the need to culture and characterise large numbers of individual isolates.

## Materials and methods

### Nodule sample collection

White clover (*Trifolium repens*) root nodules were sampled from three clover field trials under identical management in Denmark (Store Heddinge, DK), France (Rennes, F), and the United Kingdom (Didbrook, UK). At each site, 18 F2 families (1-18) and two commercial varieties (K and C) were grown in duplicate in a similar block setup (**Table S1** and **Figure S1**). In addition, 50 samples were collected from Danish clover fields under organic management. The clover varieties grown at each field can be seen in Table S1. A total of 170 samples was collected (**Figure 1, Table S1**, UK=40, F=40, DK=40, DKO1=14, DKO2=8, DKO3=3, DKO4=15, DKO5=5, DKO6=5). Nodules were stored at -20°C until DNA extraction.

### Soil sampling and chemical analysis

Soil (400g) was collected from a representative spot in each DKO field. For clover field trials 6 representative samples of 400g soil were collected in the clover-free aisles between the plots. Each plot was then assigned the values of the nearest soil sampling site. Soil chemical analysis was performed by AGROLAB Agrar und Umwelt GmbH (Germany). Traits measured were: pH, phosphorus (mg/100g), potassium (mg/100g), magnesium (mg/100g), copper (mg/kg), humus, coarse sand (0.2-2mm), fine sand (0.02-0.2mm), silt (0.002-0.02mm), and clay (<0.002mm).

### DNA extraction, library preparation, and sequencing

Nodule samples were thawed at ambient temperature and crushed using a sterile homogeniser stick. DNA was extracted from the homogenised nodule samples using the DNeasy PowerLyzer PowerSoil DNA isolation kit (QIAGEN, USA). DNA was amplified for two core genes (*rpoB* and *recA*) and two accessory genes (*nodA* and *nodD*). Amplification, library preparation, and sequencing was done using the MAUI-seq method (Fields *et al*., 2019). Sequencing was done on a Illumina MiSeq (2×300bp paired end reads) by the University of York Technology Facility.

The amplification and library preparation reported in this study were done at an early stage of method development, leading to some missing data (**Figure 2**, nSamples_rpoB: 105, nSamples_recA: 153, nSamples_nodA: 129, nSamples_nodD: 130). The method has been improved since then and now has a robust and reliable amplification rate and sample recovery (Fields *et al*., 2019; Boivin *et al*., 2020).

### Read processing and data analysis

Paired-end reads were merged using the PEAR assembler (Zhang *et al*., 2014). Reads were separated by gene, filtered using the MAUI-seq method using a secondary/primary read ratio of 0.7 and a filter of 0.1% UMI abundance was added as previously described (Fields *et al*., 2019).

Neighbour-joining phylogenetic trees were constructed using MEGAX software with 500 bootstrap repetitions. *Rlt* reference sequences for all four genes were extracted from the 196 strains with available whole genome sequencing data (Cavassim *et al*., 2020). Relative allele abundance was calculated for both the MAUI-seq data and the 196 *Rlt* genomes. Raw UMI counts are shown in **Figure S3**. Geographical maps were generated using the R packages ‘maps’ and ‘ggplot2’ (R Core team, 2015; Wickham, 2016). Heatmaps were generated from relative allele abundance of individual genes using ‘ggplot2’. The hierarchical *F*-statistics (*F*_ST_) were calculated using the ‘varcomp.glob’ function and tested using ‘test.between’ and ‘test.within’ in the ‘Hierfstat’ R package (Goudet & Jombart, 2015). Correlations and associated p-values were calculated using the ‘agricolae’ R package (de Mendiburu, 2014). Pairwise geographic distances were calculated using the ‘geosphere’ R package (Hijmans *et al*., 2019). Mantel tests were performed using 5000 repetitions in the ‘ade4’ R package (Dray & Dufour, 2007). Correlations between soil chemical properties and allele frequency were done using base R and visualised using ‘corrplot’ (Wei & Simko, 2016).

Nucleotide diversity (π, the average number of nucleotide differences per site between two DNA sequences in all possible pairs in the sample population) was calculated for each individual sample within the DKO and field trial data using a custom script.

## Acknowledgements

We thank David Sherlock for his help with developing the method, the University of York Technology Facility for sequencing, Asger Bachmann and Terry Mun for preliminary data analysis and script development, SEGES and the farmers for access to the organic fields, and DLF for access to their clover field trials. This work was funded by grant no. 4105-00007A from Innovation Fund Denmark (S.U.A.). Initial development of the method was funded by the EU FP7-KBBE project LEGATO (J.P.W.Y).

## Author contributions

Conceptualization: JPWY; Methodology: JPWY, SM; Software: SM, MT, JPWY; Validation: SM; Formal analysis: SM, MT, JPWY; Investigation: SM; Resources: JPWY, SUA ; Data curation: SM, JPWY; Writing - original draft: SM; Writing - review and editing: SM, MT, JPWY, SUA; Visualisation: SM; Supervision: JPWY, SUA; Project administration: JPWY, SUA; Funding acquisition: JPWY, SUA.

## Data accessibility

Raw Illumina reads are available in the SRA repositories with accession number [in progress] (Moeskjær *et al*., 2020). MAUI-seq scripts are available in the GitHub repository https://github.com/jpwyoung/MAUI. A detailed protocol for sampling, sample preparation, and read processing is available in Fields et al., 2020 (Fields *et al*., 2019). Sampling locations, soil chemical data, and clover genotype data is available in **Table S1**.

## References

Andrews, M., de Meyer, S., James, E.K., Stepkowski, T., Hodge, S., Simon, M.F. & Young, J.P.W. (2018) Horizontal transfer of symbiosis genes within and between rhizobial genera: Occurrence and importance. Genes..

Baudoin, E., Benizri, E. & Guckert, A. (2003) Impact of artificial root exudates on the bacterial community structure in bulk soil and maize rhizosphere. Soil Biology and Biochemistry.

Berendsen, R.L., Pieterse, C.M.J. & Bakker, P.A.H.M. (2012) The rhizosphere microbiome and plant health. Trends in Plant Science..

Boivin, S., Ait Lahmidi, N., Sherlock, D., Bonhomme, M., Dijon, D., Heulin-Gotty, K., Le-Queré, A., Pervent, M., Tauzin, M., Carlsson, G., Jensen, E., Journet, E.P., Lopez-Bellido, R., Seidenglanz, M., Marinkovic, J., Colella, S., Brunel, B., Young, P. & Lepetit, M. (2020) Host-specific competitiveness to form nodules in Rhizobium leguminosarum symbiovar viciae. New Phytologist, 226, 555–568.

van Cauwenberghe, J., Verstraete, B., Lemaire, B., Lievens, B., Michiels, J. & Honnay, O. (2014) Population structure of root nodulating Rhizobium leguminosarum in Vicia cracca populations at local to regional geographic scales. Systematic and Applied Microbiology.

Cavassim, M.I.A., Moeskjær, S., Moslemi, C., Fields, B., Bachmann, A., Vilhjálmsson, B.J., Schierup, M.H., Young, J.P.W. & Andersen, S.U. (2020) Symbiosis genes show a unique pattern of introgression and selection within a rhizobium leguminosarum species complex. Microbial Genomics.

Coller, E., Cestaro, A., Zanzotti, R., Bertoldi, D., Pindo, M., Larger, S., Albanese, D., Mescalchin, E. & Donati, C. (2019) Microbiome of vineyard soils is shaped by geography and management. Microbiome.

Costa, R., Salles, J.F., Berg, G. & Smalla, K. (2006) Cultivation-independent analysis of Pseudomonas species in soil and in the rhizosphere of field-grown Verticillium dahliae host plants. Environmental Microbiology.

Dray, S. & Dufour, A.B. (2007) The ade4 package: Implementing the duality diagram for ecologists. Journal of Statistical Software.

Efrose, R.C., Rosu, C.M., Stedel, C., Stefan, A., Sirbu, C., Gorgan, L.D., Labrou, N.E. & Flemetakis, E. (2018) Molecular diversity and phylogeny of indigenous Rhizobium leguminosarum strains associated with Trifolium repens plants in Romania. Antonie van Leeuwenhoek, International Journal of General and Molecular Microbiology.

Ferguson, S., Major, A.S., Sullivan, J.T., Bourke, S.D., Kelly, S.J., Perry, B.J. & Ronson, C.W. (2020) Rhizobium leguminosarum bv. trifolii NodD2 Enhances Competitive Nodule Colonization in the Clover-Rhizobium Symbiosis. Applied and environmental microbiology.

Fields, B., Moeskjær, S., Friman, V.-P., Andersen, S.U. & Young, J.P.W. (2019) MAUI-seq: Multiplexed, high-throughput amplicon diversity profiling using unique molecular identifiers. bioRxiv.

Goudet, J. & Jombart, T. (2015) Estimation and Tests of Hierarchical F-Statistics. R Core Team.

Gutjahr, C. & Parniske, M. (2013) Cell and developmental biology of arbuscular mycorrhiza symbiosis. Annual Review of Cell and Developmental Biology..

Hahn, M.W., Koll, U., Jezberová, J. & Camacho, A. (2015) Global phylogeography of pelagic Polynucleobacter bacteria: Restricted geographic distribution of subgroups, isolation by distance and influence of climate. Environmental Microbiology.

Hansen, B., Kristensen, E.S., Grant, R., Høgh-Jensen, H., Simmelsgaard, S.E. & Olesen, J.E. (2000) Nitrogen leaching from conventional versus organic farming systems - A systems modelling approach. European Journal of Agronomy.

Heath, K.D. & Tiffin, P. (2009) Stabilizing mechanisms in a legume-rhizobium mutualism. Evolution.

Hijmans, R.J., Williams, E. & Vennes, C. (2019) geosphere: Spherical Trigonometry. R package version 1.5-10. package geosphere..

Hoetzinger, M., Schmidt, J., Jezberová, J., Koll, U. & Hahn, M.W. (2017) Microdiversification of a pelagic Polynucleobacter species is mainly driven by acquisition of genomic islands from a partially interspecific gene pool. Applied and Environmental Microbiology.

Kelly, S., Sullivan, J.T., Kawaharada, Y., Radutoiu, S., Ronson, C.W. & Stougaard, J. (2018) Regulation of Nod factor biosynthesis by alternative NodD proteins at distinct stages of symbiosis provides additional compatibility scrutiny. Environmental Microbiology.

Kumar, N., Lad, G., Giuntini, E., Kaye, M.E., Udomwong, P., Jannah Shamsani, N., Peter W Young, J. & Bailly, X. (2015) Bacterial genospecies that are not ecologically coherent: Population genomics of rhizobium leguminosarum 140133. Open Biology, 5.

Leff, J.W., Jones, S.E., Prober, S.M., Barberán, A., Borer, E.T., Firn, J.L., Harpole, W.S., Hobbie, S.E., Hofmockel, K.S., Knops, J.M.H., McCulley, R.L., la Pierre, K., Risch, A.C., Seabloom, E.W., Schütz, M., Steenbock, C., Stevens, C.J. & Fierer, N. (2015) Consistent responses of soil microbial communities to elevated nutrient inputs in grasslands across the globe. Proceedings of the National Academy of Sciences of the United States of America.

Martínez-Hidalgo, P. & Hirsch, A.M. (2017) The nodule microbiome: N2fixing rhizobia do not live alone. Phytobiomes Journal..

Mendes, L.W., Tsai, S.M., Navarrete, A.A., de Hollander, M., van Veen, J.A. & Kuramae, E.E. (2015) Soil-Borne Microbiome: Linking Diversity to Function. Microbial Ecology.

de Mendiburu, F. (2014) Agricolae: Statistical procedures for agricultural research. R package version 1.2-0. http://CRAN.R-project.org/package=agricolae.

Moeskjær, S., Tausen, M., Andersen, S.U. & Young, J.P.W. (2020) MiSeq of Rhizobium leguminosarum bv. trifolii: rpoB, recA, nodA and nodD amplicons from T. repens nodules from field trials and organic fields. SRA accession number [in progress].

Oldroyd, G.E.D., Murray, J.D., Poole, P.S. & Downie, J.A. (2011) The rules of engagement in the legume-rhizobial symbiosis. Annual Review of Genetics.

Palmer, K.M. & Young, J.P.W. (2000) Higher diversity of Rhizobium leguminosarum biovar viciae populations in arable soils than in grass soils. Applied and Environmental Microbiology.

Panke-Buisse, K., Poole, A.C., Goodrich, J.K., Ley, R.E. & Kao-Kniffin, J. (2015) Selection on soil microbiomes reveals reproducible impacts on plant function. ISME Journal.

R Core team. (2015) R Core Team. R: A Language and Environment for Statistical Computing. R Foundation for Statistical Computing, Vienna, Austria. ISBN 3-900051-07-0, URL http://www.R-project.org/.<http://www.mendeley.com/research/r-language-environment-statistical-computing-96/ papers2://publication/uuid/A1207DAB-22D3-4A04-82FB-D4DD5AD57C28>.

Ramette, A. & Tiedje, J.M. (2007) Multiscale responses of microbial life to spatial distance and environmental heterogeneity in a patchy ecosystem. Proceedings of the National Academy of Sciences of the United States of America.

Rosselló-Mora, R., Lucio, M., Pẽa, A., Brito-Echeverría, J., López-López, A., Valens-Vadell, M., Frommberger, M., Antón, J. & Schmitt-Kopplin, P. (2008) Metabolic evidence for biogeographic isolation of the extremophilic bacterium Salinibacter ruber. ISME Journal.

Sbabou, L., Regragui, A., Filali-Maltouf, A., Ater, M. & Béna, G. (2016) Local genetic structure and worldwide phylogenetic position of symbiotic Rhizobium leguminosarum strains associated with a traditional cultivated crop, Vicia ervilia, from Northern Morocco. Systematic and Applied Microbiology.

Schlatter, D., Kinkel, L., Thomashow, L., Weller, D. & Paulitz, T. (2017) Disease suppressive soils: New insights from the soil microbiome. Phytopathology..

Schreiter, S., Ding, G.C., Heuer, H., Neumann, G., Sandmann, M., Grosch, R., Kropf, S. & Smalla, K. (2014) Effect of the soil type on the microbiome in the rhizosphere of field-grown lettuce. Frontiers in Microbiology.

Smalla, K., Wieland, G., Buchner, A., Zock, A., Parzy, J., Kaiser, S., Roskot, N., Heuer, H. & Berg, G. (2001) Bulk and Rhizosphere Soil Bacterial Communities Studied by Denaturing Gradient Gel Electrophoresis: Plant-Dependent Enrichment and Seasonal Shifts Revealed. Applied and Environmental Microbiology.

Stajkovic-Srbinovic, O., de Meyer, S.E., Miličić, B., Delic, D. & Willems, A. (2012) Genetic diversity of rhizobia associated with alfalfa in Serbian soils. Biology and Fertility of Soils.

Stefan, A., van Cauwenberghe, J., Rosu, C.M., Stedel, C., Labrou, N.E., Flemetakis, E. & Efrose, R.C. (2018) Genetic diversity and structure of Rhizobium leguminosarum populations associated with clover plants are influenced by local environmental variables. Systematic and Applied Microbiology.

Thies, J.E., Singleton, P.W. & Bohlool, B.B. (1991) Influence of the size of indigenous rhizobial populations on establishment and symbiotic performance of introduced rhizobia on field-grown legumes. Applied and Environmental Microbiology.

Tian, C.F., Young, J.P.W., Wang, E.T., Tamimi, S.M. & Chen, W.X. (2010) Population mixing of Rhizobium leguminosarum bv. viciae nodulating Vicia faba: The role of recombination and lateral gene transfer. FEMS Microbiology Ecology.

Vos, M. & Velicer, G.J. (2008) Isolation by Distance in the Spore-Forming Soil Bacterium Myxococcus xanthus. Current Biology.

Wakelin, S., Tillard, G., van Ham, R., Ballard, R., Farquharson, E., Gerard, E., Geurts, R., Brown, M., Ridgway, H. & O’Callaghan, M. (2018) High spatial variation in population size and symbiotic performance of Rhizobium leguminosarum bv. trifolii with white clover in New Zealand pasture soils. PLoS ONE.

Wei, T. & Simko, V. (2016) The corrplot package. R Core Team.

Whitaker, R.J., Grogan, D.W. & Taylor, J.W. (2003) Geographic barriers isolate endemic populations of hyperthermophilic archaea. Science.

Wickham, H. (2016) Ggplot2. P. in: Ggplot2: Elegant Graphics for Data Analysis. Springer-Verlag New York.

Zhang, J., Kobert, K., Flouri, T. & Stamatakis, A. (2014) PEAR: A fast and accurate Illumina Paired-End reAd mergeR. Bioinformatics, 30, 614–620.

Zhao, Z.B., He, J.Z., Geisen, S., Han, L.L., Wang, J.T., Shen, J.P., Wei, W.X., Fang, Y.T., Li, P.P. & Zhang, L.M. (2019) Protist communities are more sensitive to nitrogen fertilization than other microorganisms in diverse agricultural soils. Microbiome.

