## Supplementary figures for "Amplicons and isolates: *Rhizobium* diversity in fields under conventional and organic management"

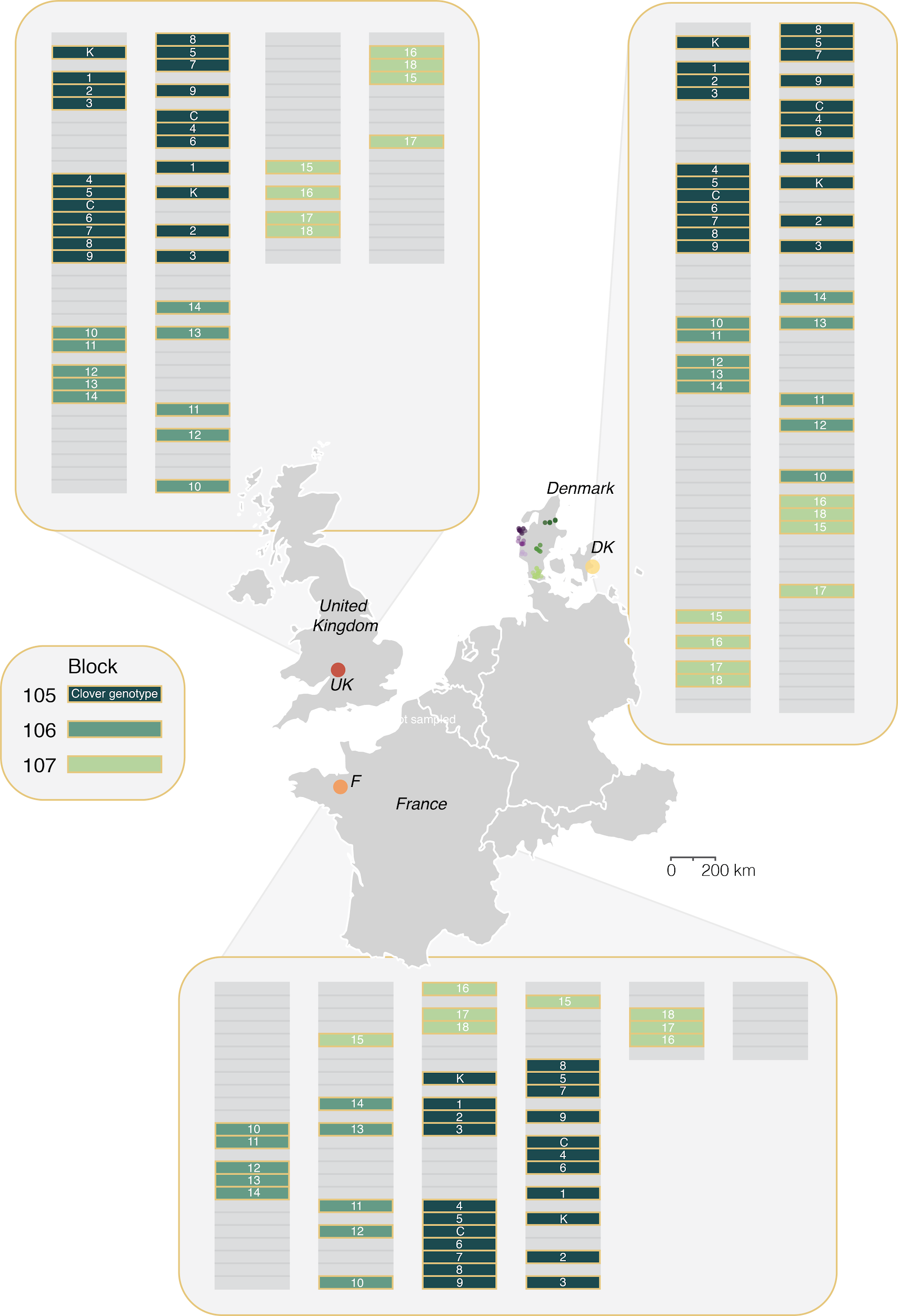

**Figure S1.** Plot layout for the field trial sites in Store Heddinge, Denmark (DK), Rennes, France (F), and Didbrook, United Kingdom (UK). The trial was set up in three blocks (105, 106, and 107). Block layout was replicated between trial sites. The same F2 clover families (1-18) were grown at each site, along with two commercial varieties (K and C, **Table S1**). All clover families/varieties were sampled in duplicate within each trial site.

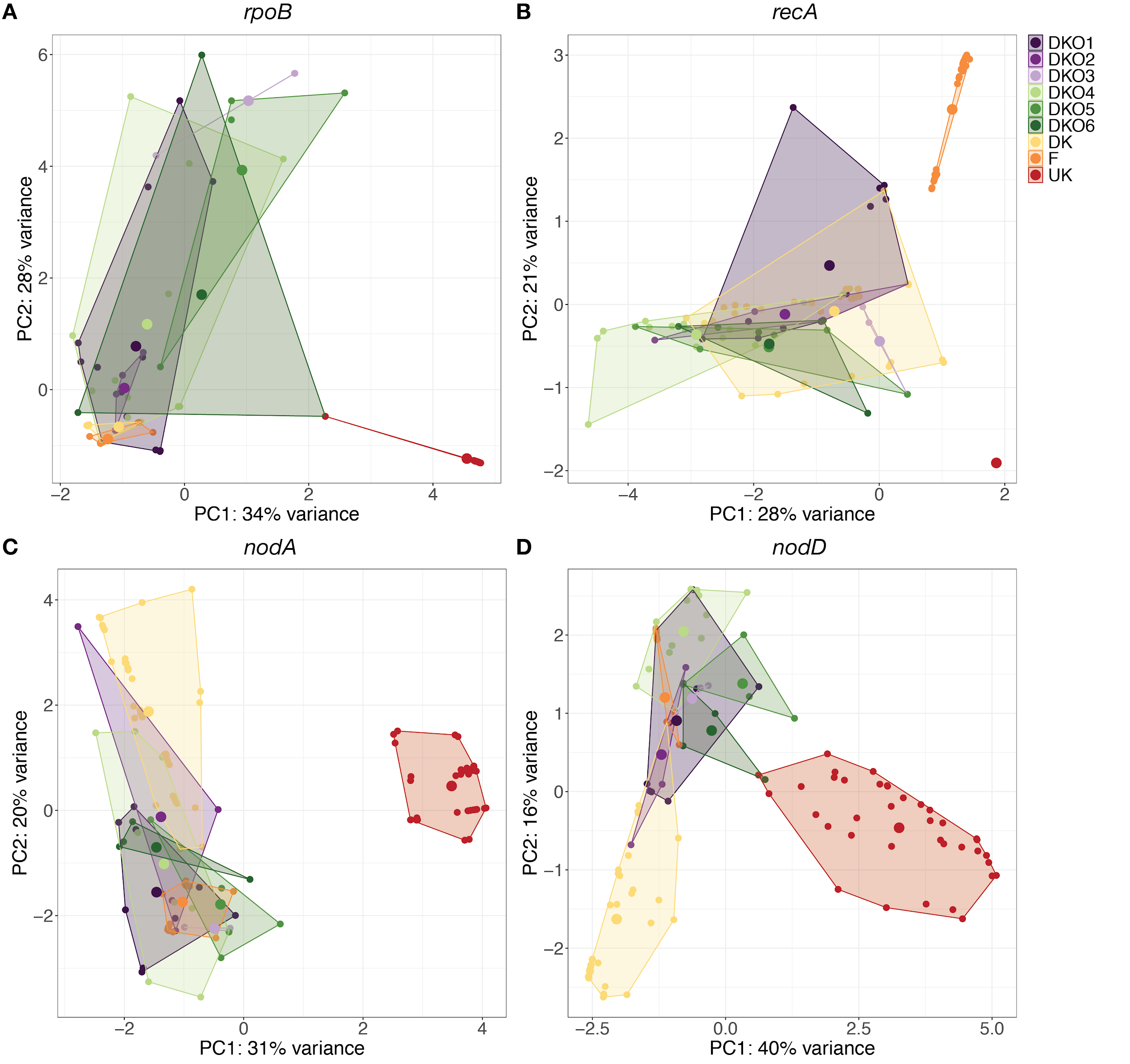

**Figure S2.** Individual Principal Components Analysis of four *Rhizobium leguminosarum* symbiovar *trifolii* genes clustered by MAUI-seq. DNA was isolated from 100 nodules for each point. **A:** *rpoB,* **B:** *recA,* **C:** *nodA* and **D:** *nodD*.

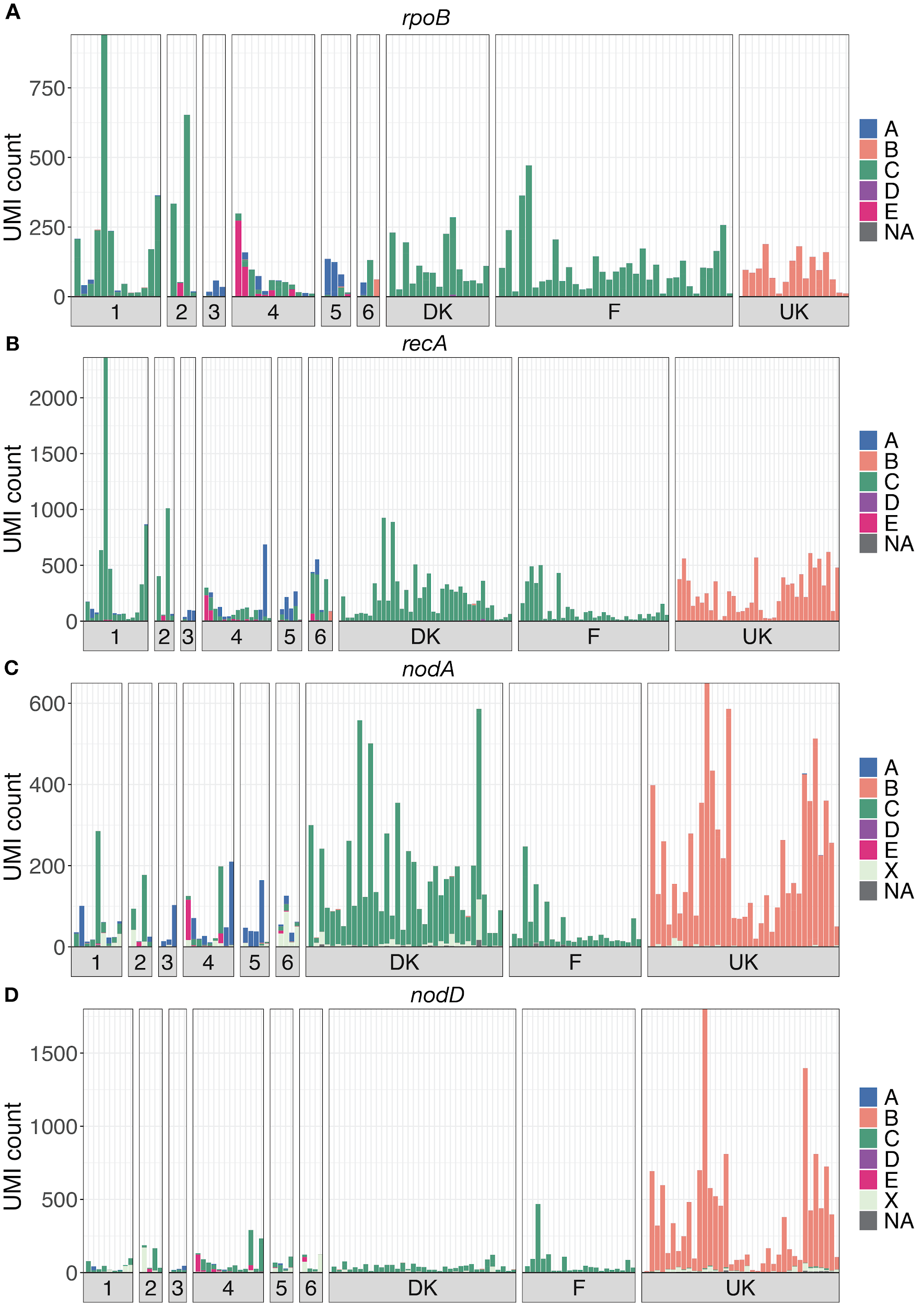

**Figure S3.** Genospecies composition of each individual sample per gene displayed by the raw UMI count. The DKO groupings are labelled by their respective number (DKO1=1). Core genes (*rpoB* and *recA*) are assigned to the genospecies A-E (Kumar et al., 2015). Nod genes (*nodA* and *nodD*) are assigned to a genospecies if possible, or to a clade of introgressing genes labelled X (Cavassim et al., 2020). If an amplicon could not clearly be assigned to a clade it is marked as NA.

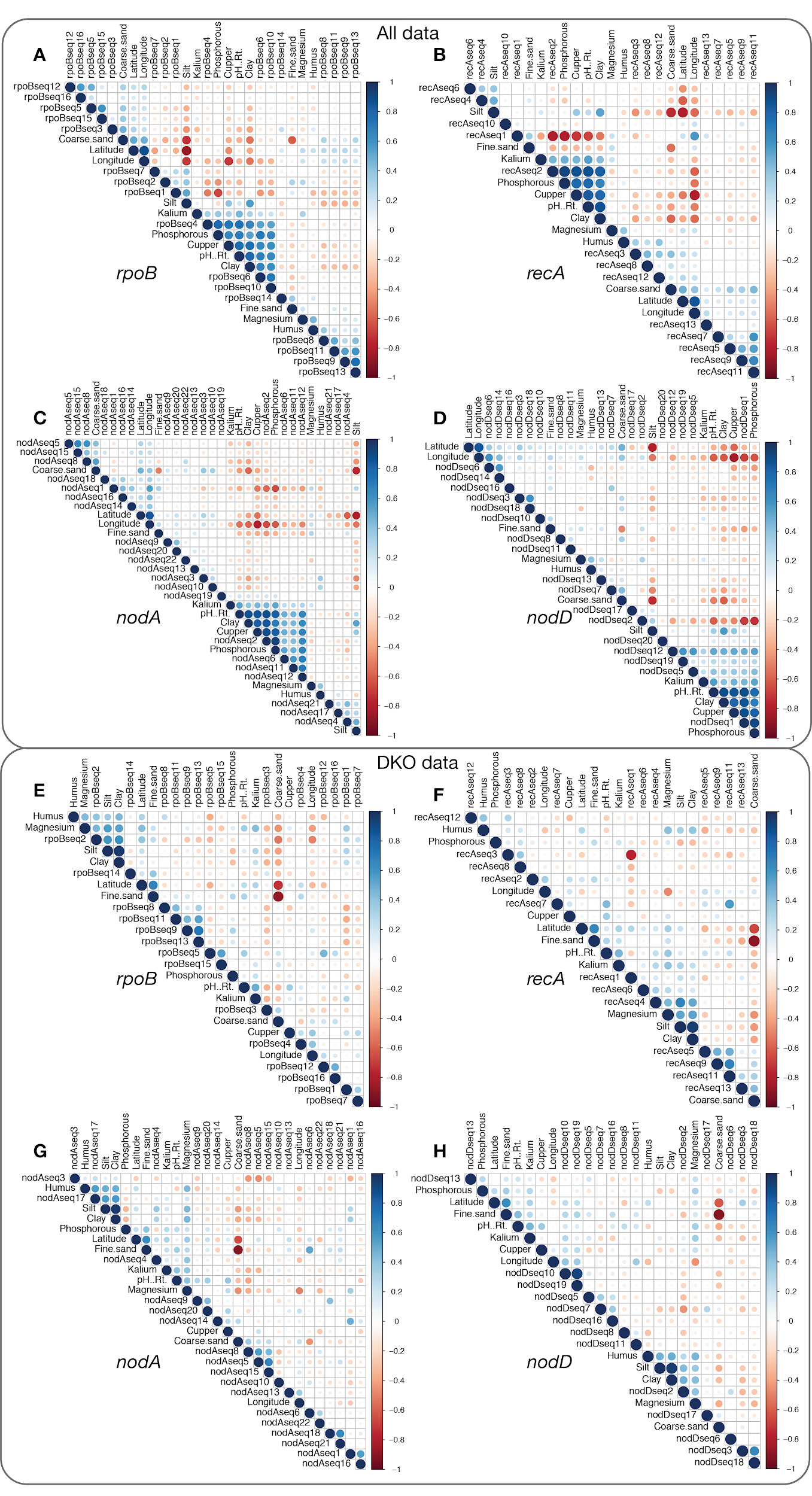

**Figure S4**. Correlations between normalised allele frequencies and soil chemical properties per gene for **A-D**: the full dataset (All data) and **E-H**: the DKO subset. Panel **E** and **H** are also shown in **Figure 4** in the main document.

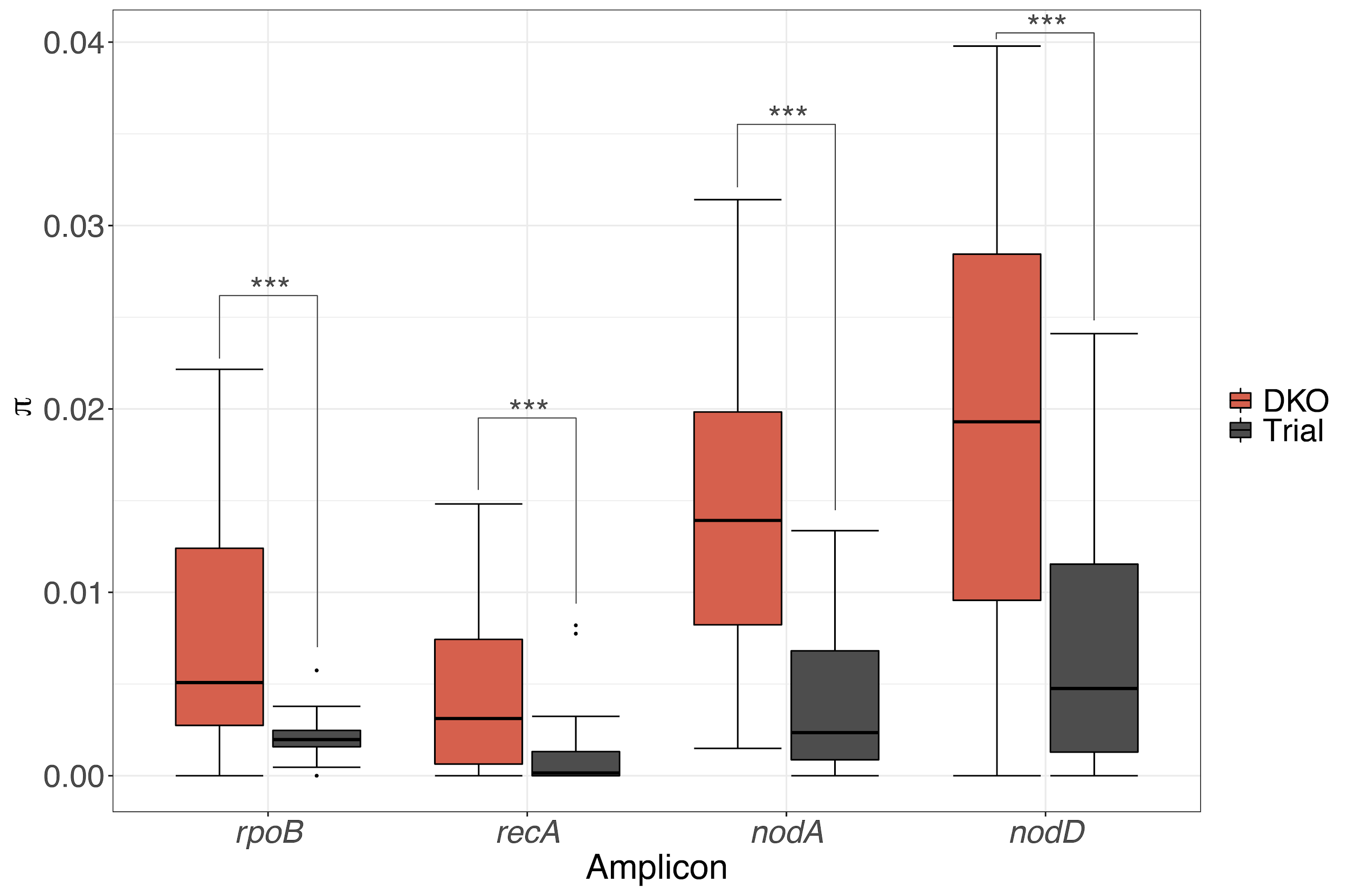

**Figure S5**. Nucleotide diversity within populations. Average number of nucleotide differences per site between two DNA sequences in all possible pairs in the sample population (𝛑) for each individual sample within the DKO and field trial data. Dots illustrate the **𝛑** value for each individual sample. Bars represent the first and third quartiles, with the solid line denoting the median. Whiskers correspond to the 1.5 * interquartile range. p-values were calculated using Welch Two Sample unequal variances t-test between DKO and clover field trial samples and are indicated by asterisks; *p<0.05, **p<0.01, and ***p<0.001. Sample sizes for each gene; *rpoB*: nDKO=39 and nTrial=66 (p-value=3.007e-06), *recA*: nDKO=46 and nTrial=107 (p-value=8.601e-07), *nodA*: nDKO=34 and nTrial=95 (p-value=1.916e-09), and *nodD*: nDKO=37 and nTrial=93 (p-value=2.875e-08).
